# Classification and Mutation Prediction from Non-Small Cell Lung Cancer Histopathology Images using Deep Learning

**DOI:** 10.1101/197574

**Authors:** Nicolas Coudray, Andre L. Moreira, Theodore Sakellaropoulos, David Fenyö, Narges Razavian, Aristotelis Tsirigos

## Abstract

Visual analysis of histopathology slides of lung cell tissues is one of the main methods used by pathologists to assess the stage, types and sub-types of lung cancers. Adenocarcinoma and squamous cell carcinoma are two most prevalent sub-types of lung cancer, but their distinction can be challenging and time-consuming even for the expert eye. In this study, we trained a deep learning convolutional neural network (CNN) model (inception v3) on histopathology images obtained from The Cancer Genome Atlas (TCGA) to accurately classify whole-slide pathology images into adenocarcinoma, squamous cell carcinoma or normal lung tissue. Our method slightly outperforms a human pathologist, achieving better sensitivity and specificity, with ∼0.97 average Area Under the Curve (AUC) on a held-out population of whole-slide scans. Furthermore, we trained the neural network to predict the ten most commonly mutated genes in lung adenocarcinoma. We found that six of these genes – STK11, EGFR, FAT1, SETBP1, KRAS and TP53 – can be predicted from pathology images with an accuracy ranging from 0.733 to 0.856, as measured by the AUC on the held-out population. These findings suggest that deep learning models can offer both specialists and patients a fast, accurate and inexpensive detection of cancer types or gene mutations, and thus have a significant impact on cancer treatment.

## Introduction

According to the American Cancer Society^1^, over 150,000 lung cancer patients succumb to their disease each year, while another 200,000 new cases are diagnosed on a yearly basis. It is one of the most widely spread cancers in the world, due mostly to smoking, but also exposure to toxic chemicals like radon, asbestos and arsenic. Non-small cell lung cancers represent 85% of the cases and three sub-types are distinguished: Adenocarcinoma (LUAD), Squamous Cell carcinoma (LUSC) and, most rarely, large-cell carcinoma. Lung biopsies are typically used to diagnose lung cancer subtype and stage. Targeted therapies are applied depending on the type of cancer, stage and the presence of sensitizing mutations^1,2^. For example, EGFR (epidermal growth factor receptor) mutations, present in about 20% of LUAD, and ALK mutations (anaplastic lymphoma receptor tyrosine kinase), present in less than 5% of LUAD^3^, both have currently targeted therapies approved by the Food and Drug Administration (FDA)^4^. Mutations in other genes, such as KRAS and TP53 are very common (about 25% and 50% respectively), but have proven particularly challenging drug-targets so far^3,5^.

Virtual microscopy of stained images of tissues are typically acquired at magnifications of x20 to x40, generating very large two-dimensional images (10,000 to over 100,000 pixels in each dimension) that can be tricky to visually analyze in an exhaustive way. Furthermore, accurate interpretation can be difficult and the distinction between LUAD and LUSC is not always clear, particularly in poorly-differentiated tumors, where ancillary studies is recommended for accurate classification. To assist experts, automatic analysis of lung cancer whole-slide images has been recently studied for survival prognosis^6^ and classification^7^. In these studies, Yu et al. combined conventional thresholding and image processing techniques with machine learning methods, such as random forest classifiers, SVM or Naïve Bayes classifiers, achieving an Area Under the Curve (AUC) of ∼0.85 in distinguishing normal from tumor slides, and ∼0.75 in distinguishing LUAD from LUSC slides. Here, we demonstrate how the field can greatly benefit from deep learning, by presenting a strategy based on Convolutional Neural Networks (CNNs) that not only outperforms previously published work, but also achieves accuracies that are at least comparable, if not superior, to human pathologists. The development of new inexpensive and more powerful technologies with higher computing power (in particular Graphics Processing Units, GPUs) has made possible the training of larger and more complex systems^8–10^. This resulted in the design of several deep CNNs, capable of accomplishing complex visual recognition tasks. Such algorithms have already been successfully used for segmentation^11^ or classification of medical images^12^ and cancers such as breast^13–15^, colon cancers^16^ or osterosarcoma^17^. CNNs have also been studied for classifying lung patterns on CT (Computerized Tomography) scans, achieving a f-score of ∼85.5%^18^. Here, to study the automatic classification of lung cancer tissues, we used the inception v3 architecture^19^ and whole-slide images of hematoxylin and eosin stained histopathology images from TCGA obtained by excision. In 2014, Google won the ImageNet Large-Scale Visual Recognition Challenge by developing the GoogleNet architecture^20^, derived from the work from Lin et. al^21^, which increased the robustness to translation and non-linear learning abilities by using multi-layer perceptrons and global averaging pooling. Inception architecture is particularly useful for processing the data in multiple resolutions, a feature that makes this architecture suitable for pathology tasks. This network has already been successfully adapted to other specific types of classifications like skin cancers^22^ and diabetic retinopathy detection^23^.

## Results

We are here comparing several approaches for the classification of tumor slides. First, we employed a strategy similar to the one used by Yu et al ^7^, consisting of a two-step binary classification of normal versus tumor slides, followed by a classification of LUAD versus LUSC slides. We then explored a direct classification of the three types of whole-slide images. Finally, we further analyzed LUAD slides to identify which gene mutations could be predicted from those images: we modified and trained the inception v3 architecture on the 10 most common mutations found in the TCGA dataset and related to lung cancer. In this study, we also compare two training approaches: transfer learning versus fully training the inception architecture. In the first case, most of the network keeps the weights learned after the network was trained on object recognition task on the ImageNet dataset, while only the last layer (fully connected layer) of the network is trained. In the second case, the weights are reinitialized randomly, and the network is trained end-to-end, using exclusively lung cancer images.

### Fully-trained inception v3 network provides accurate diagnosis (AUC=0.97) of lung histopathology images

The TCGA dataset characteristics and our overall computational strategy are summarized in **Figure 1** (see method section for details). We used 1634 whole-slide images from the Genomic Data Commons database: 1176 tumor tissues and 459 normal (**Figure 1a).** These whole-slide images were split into three sets: training, validation and testing (**Figure 1d**). Because the sizes of the whole-slide images are too large to be used as direct input to a neural network (sometimes over 100,000 pixels wide, **Figure 1b**), the network was instead trained, validated and tested using 512x512 pixel tiles, obtained from non-overlapping windows of the whole-slide images. This resulted in tens to thousands of tiles per slide depending on the original size (**Figure 1c**). These tiles were first processed individually by the network, and then, per-slide aggregation (see Methods for details) of the results generated a diagnosis for each slide.

**Figure 1.**
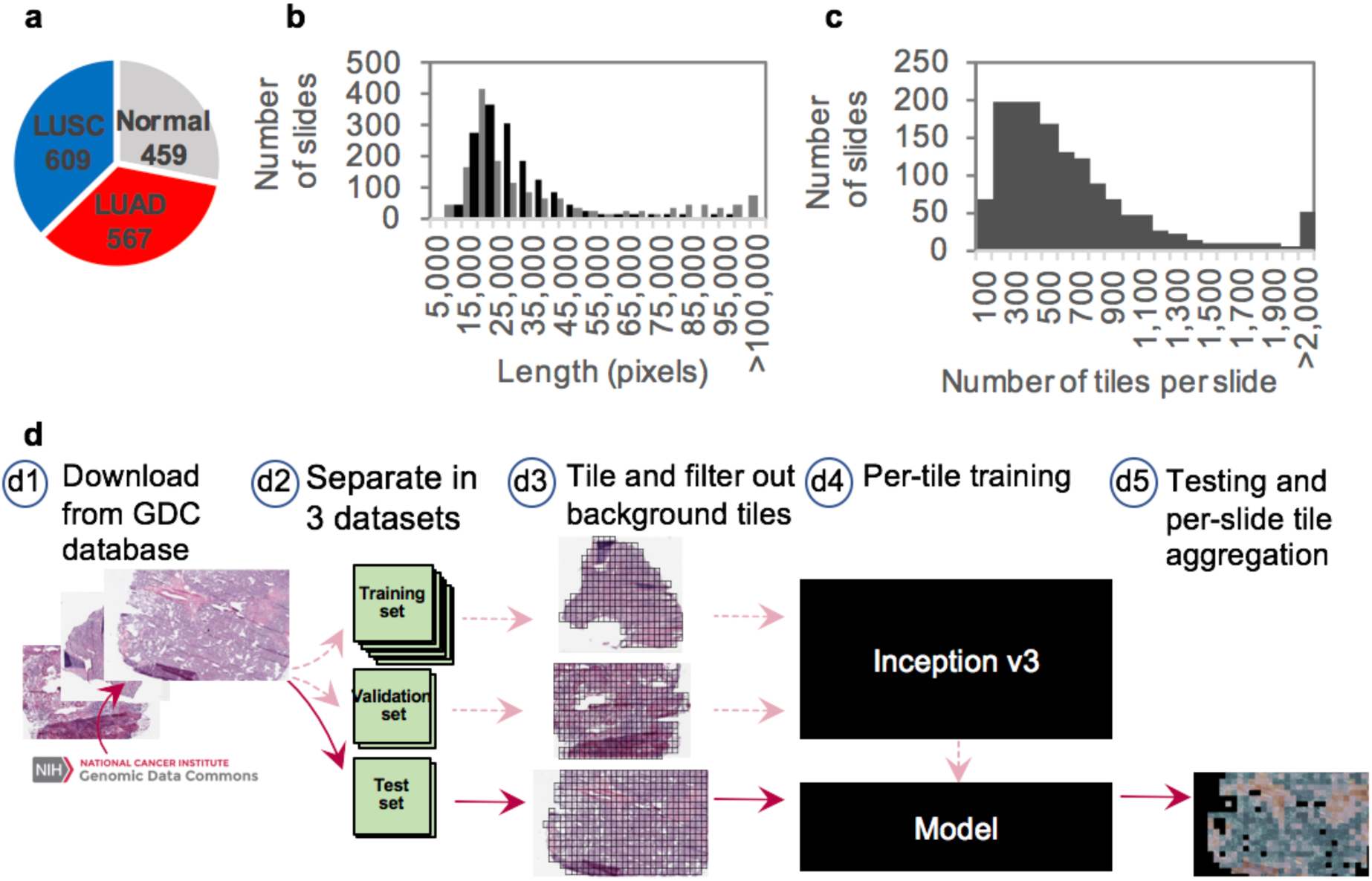
Data and strategy: **(a)** Number of whole-slide images per class. **(b)** Size distribution of the images widths (gray) and heights (black). **(c)** Distribution of the number of tiles per slide.**(d)** Strategy: **(d1)** Images of lung cancer tissues were first downloaded from the Genomic Data Common database; **(d2)** slides were then separated into a training (70%), a validation (15%) and a test set (15%); **(d3)** slides were tiled by non-overlapping 512x512 pixels windows, omitting those with over 50% background; **(d4)** the Inception v3 architecture was used and partially or fully retrained using the training and validation tiles; **(d5)** classifications were run on tiles from an independent test set and the results were finally aggregated per slide to extract the heat-maps and the AUC statistics.

Our deep learning approach effectively distinguishes tumor from normal tissue, resulting in a 96.1% per tile accuracy. To assess the accuracy on the test set, the per-tile results were aggregated on a per-slide basis either by averaging the probabilities obtained on each tile, or by counting the percentage of tiles positively classified (**Figure 2a**). This process resulted in an Area Under the Curve (AUC) of 0.990 and 0.993 (**Table 1**) respectively, outperforming the AUC of∼0.85 achieved by the feature-based approach of Yu et al.^7^. Next, we tested the performance of our approach on the more challenging task of distinguishing LUAD and LUSC. First, we testedwhether convolutional neural networks can outperform the published feature-based approach, even when plain transfer learning is used. For this purpose, the weights of the last layer of inception v3 – previously trained on the ImageNet dataset to identify 1,000 different classes – were initialized randomly and then trained for our classification task. After aggregating the statistics on a per slide basis (**Figure 2b**), this process resulted in an Area Under the Curve (AUC) of 0.847 (**Table 1**), i.e. a gain of ∼0.1 in AUC compared to the best results obtained by Yu et al^7^. using image features combined with random forest classifier^7^. The performance can be furtherimproved by fully training inception v3 leading to AUC of 0.947 when aggregation is done by computing the percentage of tiles positively classified, and to AUC of 0.950 when the aggregation is done by averaging the per-tile probabilities (**Figure 2c**). These AUC values are improved by another 0.002 when the tiles previously classified as “normal” by the first classifier are not included in the aggregation process (**Table 1** and **Figure 2d**). The ROC of such a classifier shows performance better than that of a specialist who was asked to classify the images in the test set, independently of the classification provided in TCGA (**Figure 2d**, red cross). About a third of the slides wrongly classified by the algorithm were also misclassified by the specialist, while 85% of those incorrectly classified by the specialist were properly classified by the algorithm (**Figure 2e**).

**Figure 2.**
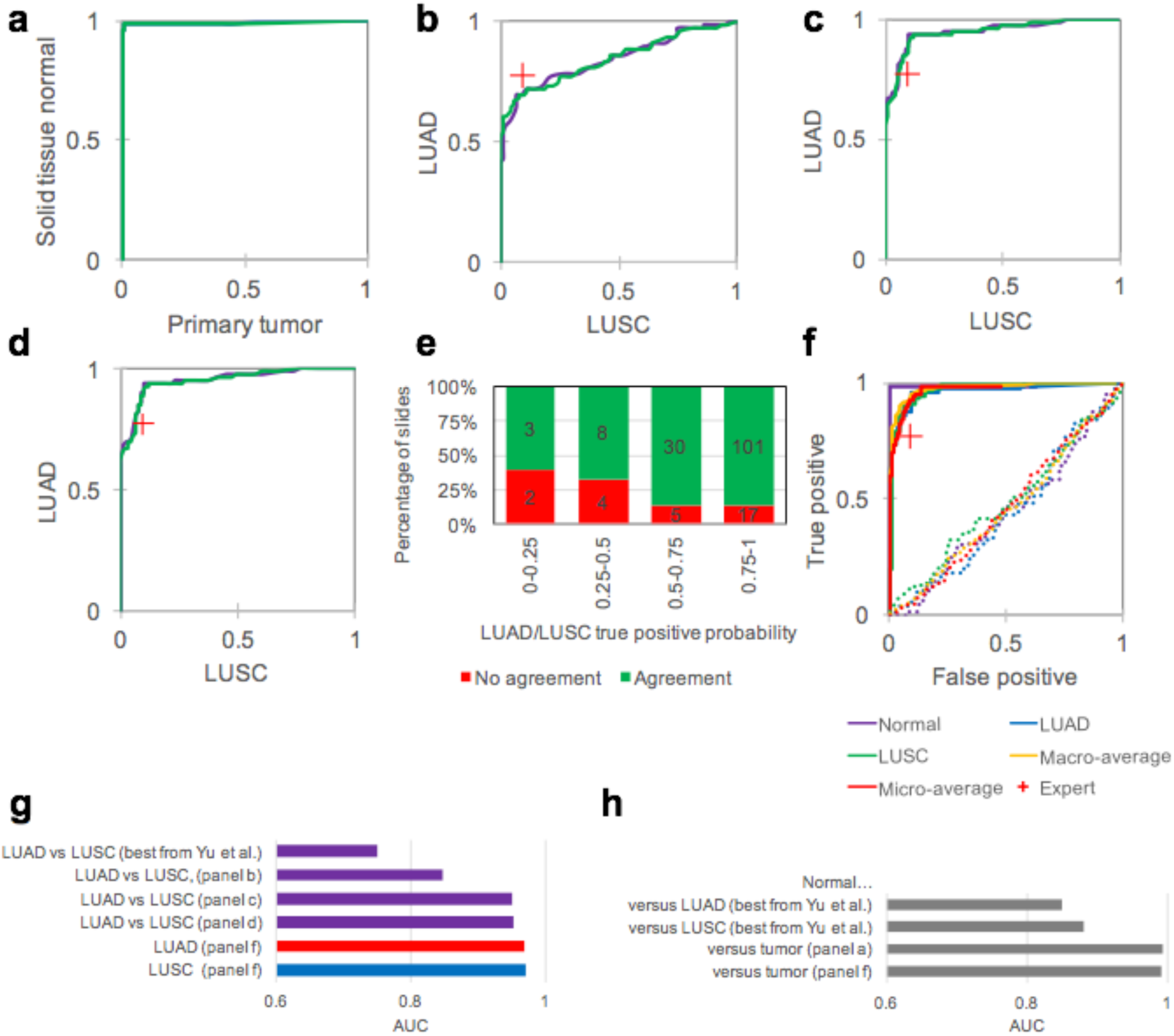
Accurate classification of lung cancer histopathology images: **(a)** Per-slide Receiver Operating Characteristic (ROC) curves after classification of normal versus tumor images resulted in an almost error-free classification. Aggregation was either done by averaging the probability scores (purple ROC curves in a to d) or by counting the percentage of properly classified tiles (green ROC curves in a to d). **(b)** The ROC curves obtained after transfer learning for LUAD vs LUSC images classification shows poorer results than when **(c)** the same network has been fully trained. The red crosses correspond to the manual classification by a specialist. **(d)** The ROC curves from (c) is only slightly improved once the tiles classified initially as “normal” have been removed. **(e)** Proportion of slides misclassified by the specialist as a function of the true positive probability assigned in (d). The number of slides are indicated on the bars. **(f)** Multi-class ROC of the Normal vs LUAD vs LUSC classification shows the best result for overall classification of cancer type. Dotted lines are negative control trained and tested after random label assignments. **(g)** Comparison of AUCs obtained with different techniques for classification of cancer type and **(h)** of normal slides.

**Figure 3a-r** show heatmap examples for LUAD and LUSC, comparing transfer-learning results with the fully trained network. In the second case, more tiles are true positive and the distribution is more homogeneous, showing for LUSC that almost all of the tiles display LUSC-like features, while for the LUAD, two regions are more prominent with LUAD-like features (one horizontal at the top, one vertical on the left) and some patches showing lower probabilities. Interestingly, most of the LUSC tiles were previously classified as tumor by the first classifier (**Figure 3t**) while for LUAD, the regions with patches having probability near 0.5 in the LUAD/LUSC classification are also those classified as normal with higher probability by the first classifier (**Figure 3s**). We investigated further the use of the deep-learning model by training and testing the network for a direct classification of the three types of images (Normal, LUAD, LUSC in **Figure 2f**). Such an approach resulted in the highest performance with all the AUCs improved to at least 0.968 (**Table 1**). **Figure 4** shows how the heat-maps are affected by such an approach: the LUSC image shows most of its tiles with a strong true positive probability of LUSC while in the LUAD image, some regions have strong LUAD features, with normal cells on the side (as confirmed by a specialist), and some light blue tiles where LUSC probability is slightly leading.

**Table 1.**
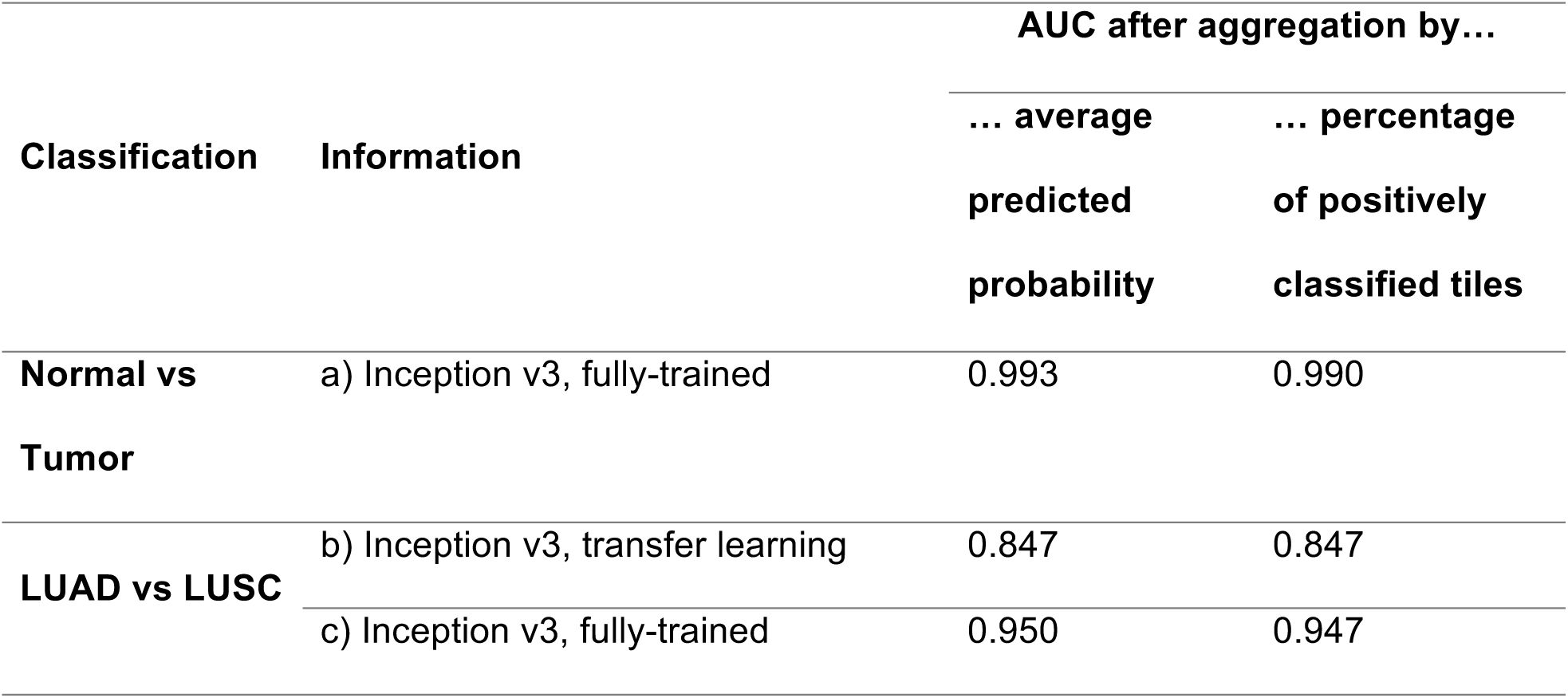
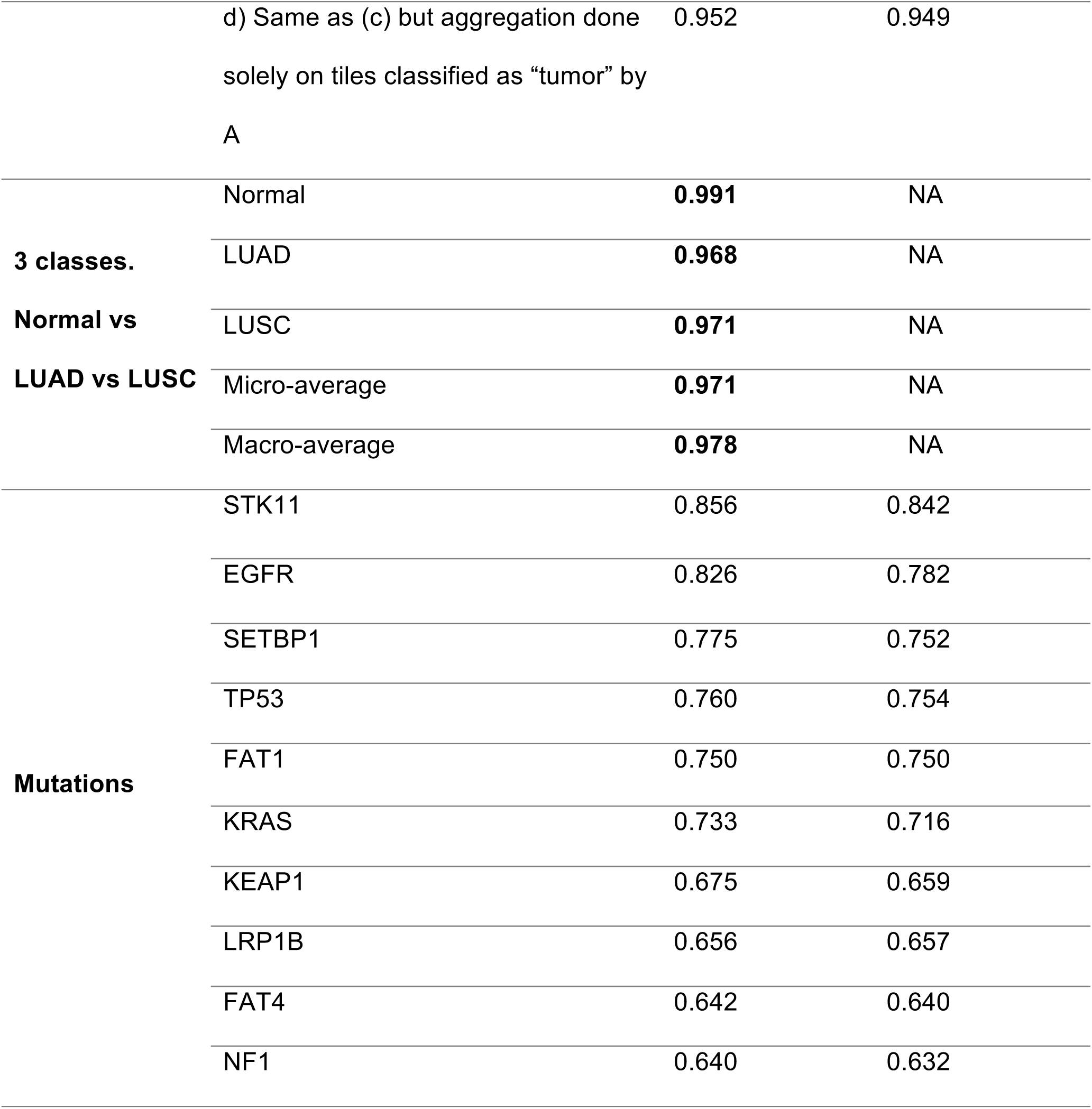
Area Under the Curve (AUC) achieved by the different classifiers

**Figure 3.**
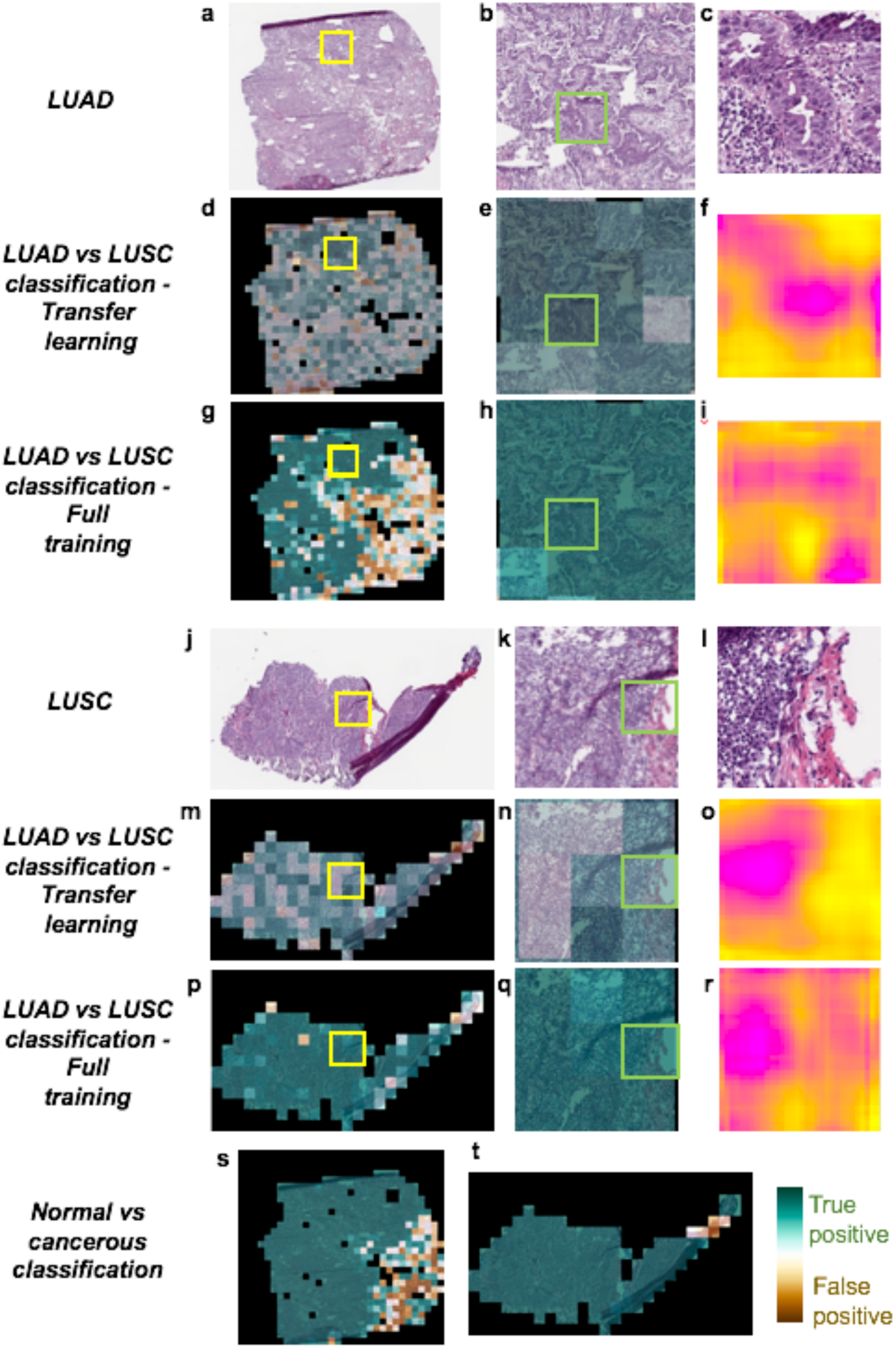
Examples of heatmaps for different binary classifications strategies: **(a)** Typical slide of Lung Adenocarcinoma (LUAD) tissue. **(b)** Zoom region corresponding to the yellow box in (a). **(c)** Tile corresponding to the green box in (b). **(d)** and **(e)** are the heat-maps corresponding to images (a) and (b), with probability assigned to each tile from brown (false positive) to green (true positive). **(f)** Per-tile heat-map generated after having applied a rolling mask on part of the tile. Yellow pixels show regions not affected by masking while the pink pixels show regions where features were important for proper classification. Images **(d)** to **(f)** were obtained after transfer learning while images **(g)** to **(i)** were obtained after fully training inception v3. Images **(j)** to **(r)** show similar examples for a Lung Squamous Cell (LUSC) tissues. **(s)** and **(t)** show the results of the “normal vs tumor” tiles classifier.

**Figure 4.**
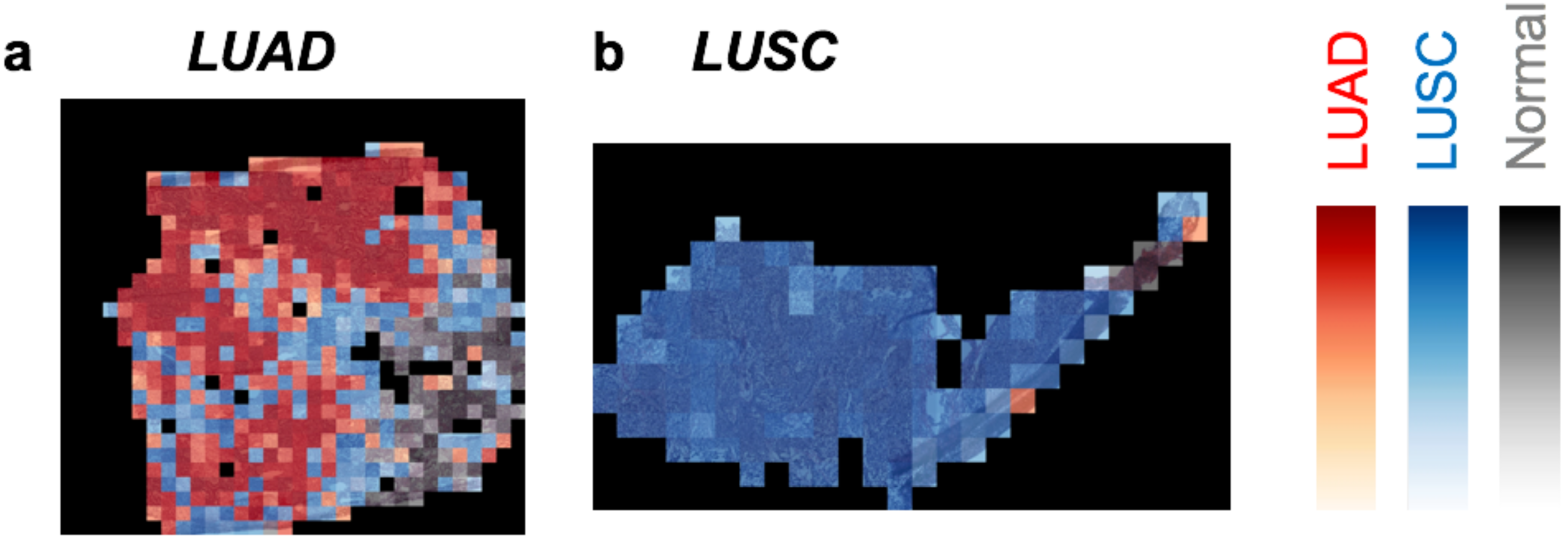
Heatmaps for classification of Normal vs LUAD vs LUSC: **(a)** and **(b)** Heatmaps corresponding to images of (**Figure 3a**) and (**Figure 3b**) with probabilities of the winning class assigned to each tile such as: red for tiles classified as LUAD, blue for LUSC and grey for Normal.

### Whole-slide images can predict 6 mutations with an AUC above 0.74

The LUAD whole-slide images were further trained to predict gene mutations. Inception v3 was modified to allow multi-output classification and tests were conducted using around 44,000 tiles from 62 slides. Box plot and ROC curves analysis (**Figure 5a-b** and **Figure supp 1**) show that at least six frequently mutated genes seem predictable using our deep learning approach: AUC values for STK11, EGFR, FAT1, SETBP1, KRAS and TP53 were found between 0.733 and 0.856 (**Table 1**). As mentioned earlier, EGFR already has targeted therapies. STK11 (Serine/Threonine protein Kinase 11), also known as Liver Kinase 1 (LKB1), is a tumor suppressor inactivated in 15-30% of non-small cell lung cancers^24^ and is also a potential therapeutic target: it has been shown on mice that phenformin, a mitochondrial inhibitor, increases survival^25^. Also, it has been shown that STK11 mutations may play a role in KRAS mutations which, combined, result in more aggressive tumors^26^. FAT1 is an ortholog of the Drosophila fat gene involved in many types of cancers and its inactivation is suspected to increase cancer cell growth^27^. Mutation of the tumor suppressor gene TP53 is thought to be more resistant to chemotherapy leading to lower survival rates in small-cell lung cancers^28^. As for SETBP1 (SET 1 binding protein), like KEAP1 and STK11, has been identified as one of the signature mutations of LUAD^29^. Finally, for each gene, we compare the classification achieved by the deep learning approach with the allele frequency (**Figure 5c**). Among the gene mutations predicted with a high AUC, four of them seem to show probabilities related to the allele frequency: FAT1, KRAS, SETBP1 and STK11.

**Figure 5.**
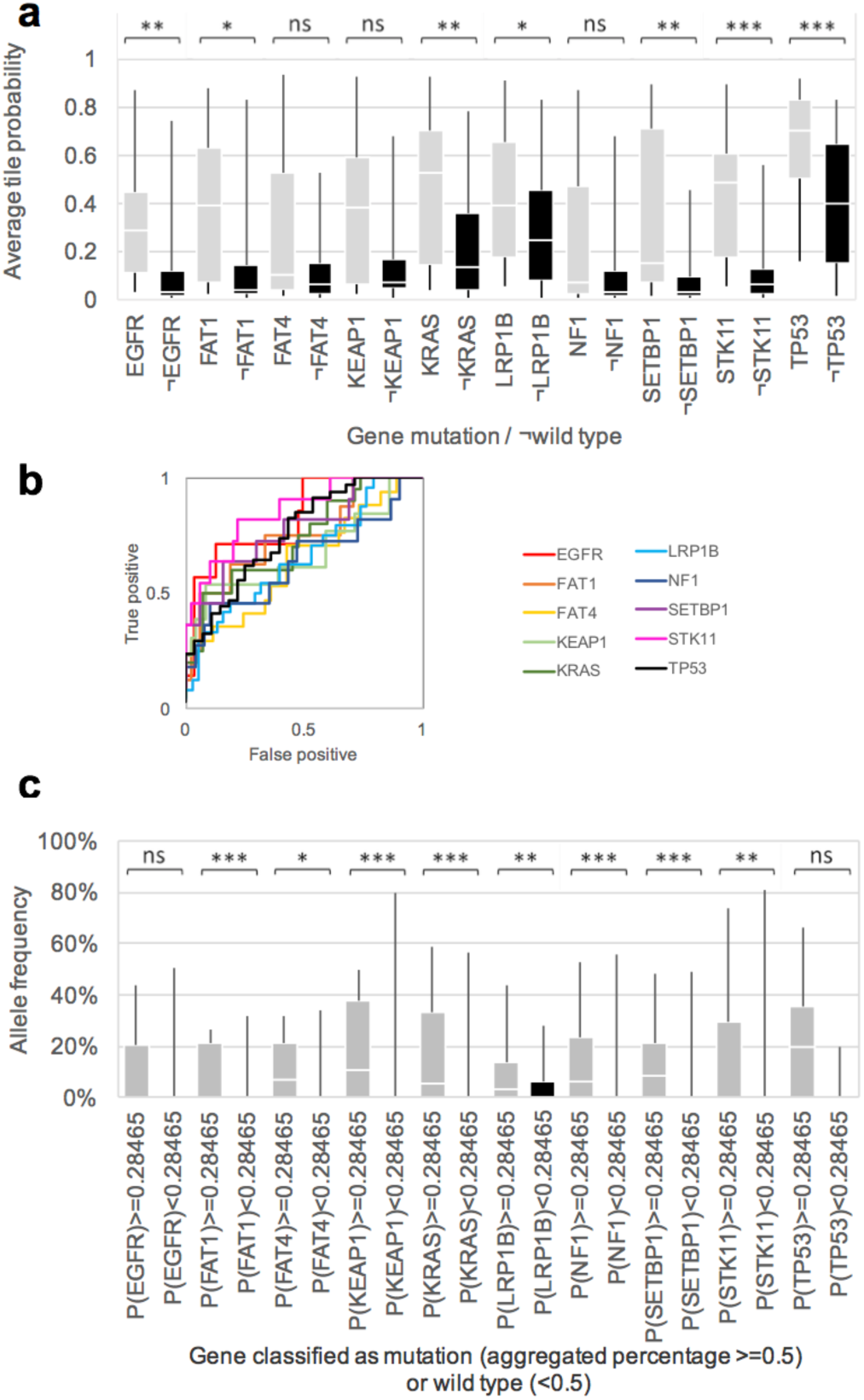
Gene mutation prediction from histopathology slides give promising results for at least 6 genes: **(a)** Mutation probability distribution for slides where each mutation is present and absent after tile aggregation done by averaging output probability. **(b)** ROC curves associated with (a). **(c)** Allele frequency as a function of slides classified by the deep-learning network as having a certain gene mutation (P≥0.5), or the wild-type (P<0.5). p-values estimated with Mann-Whitney U-test are shown as ns (p>0.05), * (p≤0.05), ** (p≤0.01) or *** (p≤0.001).

## Discussion

The analysis of lung cancer slide images using the inception v3 convolutional neural network shows a clear improvement over classification techniques combining random forest classifiers, SVM or Naïve Bayes classifiers with conventional image processing tools^7^ (**Figure 2g-h**). For LUAD vs LUSC classification, while transfer learning outperforms this previous approach by about 10% and another ∼10% is gained by fully training the network, at the expense of a much longer training period. Finally, another ∼2.8% is gained on cancer type classification when “normal” tissues are immediately considered and binary classification is replaced by a direct three-class analysis. This latest approach results in performances slightly better than those achieved by a specialist. It is interesting to notice that around a third of the slides misclassified by the algorithms have also been misclassified by the specialist, showing the intrinsic difficulty to distinguish LUAD from LUSC in some cases. However, 22 out of 26 of the slides misclassified by the specialist were assigned to the correct cancer type by the algorithm showing that it could be beneficial in assisting the specialist in their prognosis. As for classification of normal versus tumor cells, the classification is nearly unambiguous with CNN. Per-slide heat-maps (**Figure 3**) show that true positive tiles appear with a stronger probability when the network has been fully trained. For the LUAD sample, it also shows more consistency with tiles in the same adjacent regions being assigned similar probabilities while the bottom right side of the slide seems to contain less LUAD-like tumor cells according to the classification (**Figure 3g**) and is consistent with visual inspection of that region by a specialist. An example of the important features used for classification of individual is shown for LUAD (**Figure 3f,i**) and LUSC (**Figure 3o,r**). In both cases, the per-tile heat-map of the fully trained network shows a more varied gradient of colors while the tests done after transfer-learning shows more of a binary-like heat-map with regions either very yellow or very pink. The development of appropriate tools for visualizing deep learning models will help in the future to better understand the features used by the classifier^30^. In the current strategy, the only selection used for early tile removal is to make sure that there are enough information and the portion of background present is low. Afterwards, all the remaining tiles belonging to a given slide are used for training and all are associated with the label associated with it. This assumption gives good result since AUCs of 0.95 to 0.97 performance is achieved for LUSC vs LUAD, but it is unlikely that 100% of the tiles are indeed representative of the labelled cancer type. Oftentimes, the tumor is only local and some regions of the slides are not affected by the tumor. Performing an initial classification of “normal” vs “tumor” partially addresses this issue removing normal-like tiles. The fact that these are excisions of lung cancer, the tumor cells spread over the whole slide images available and not a small portion of which has clearly been beneficial for this classification. Finally, it is surprising to note the high AUCs achieved considering that several slides present artifacts inherent to freezing techniques used to prepare those samples. However, it should be noted that the available images may not fully represent the diversity that specialists have to deal with and it may be interesting in the future to assess how the network performs under the less than ideal circumstances that can occur (poor staining quality, focus not optimal or autofocus failure, lack of homogeneity in the illumination, etc). Before this study, it was a priori unclear if and how a given gene mutation would affect the pattern of tumor cells on a whole-slide image but the training of the network using mutated genes as a label lead to very promising prediction results for 6 genes: EGFR, STK11, FAT1, SETBP1, KRAS and TP53. STK11 mutation leads to the highest prediction rate with AUC above 0.85 using aggregation by averaging tile probabilities. Though the number of cases is low (44,000 tiles from 62 test slides), it is interesting to see that our training protocol gives non-random values for several genes, showing that mutation of these particular genes could be predicted from whole-slide images. Hopefully, these predictions will be confirmed once more data are made available. It means that those mutations somehow affect the way the tumor cells look like. Future work on deep-leaning models visualization tools ^30^ would help identifying those features. These probabilities could be reflecting the percentage of cells effectively affected by the mutation, the allele frequency being significantly higher for at least 4 genes when they were predicted as mutated by the neural network (**Figure 5c**). Looking, for example, at the predictions done on the whole-slide image from **Figure 3a**, our process successfully identifies TP53 (allele frequency of 0.33) and STK11 (allele frequency of 0.33) are two gene most likely mutated (**Figure supp 2a**). The heatmap shows that almost all the LUAD tiles are highly predicted as showing TP53-mutatant-like features (**Figure supp 2b**), and two major regions with STK11-mutatant-like features (**Figure supp 2c**). Interestingly, when the classification is applied on all tiles, it shows that even tiles classified as LUSC present TP53 mutations (**Figure supp 2d**) while the STK11 mutant is confined to the LUAD tiles (**Figure supp 2e**). These results are realistic since, as mentioned earlier, STK11 is a signature mutations of LUAD ^29^ while TP53 is more common in human cancers. Being able to predict gene mutations at this stage could be beneficial regarding the importance and impact of those mutations^4,24–29^. This study shows that using deep-learning convolutional neural networks for cancer analysis greatly improve the state-of-the-art automatic classification and could be a very promising tool to assist the specialist in their classification of whole-slide images of lung tissues. Histopathology slides are very large, they usually contain artifacts and be noisy with features of cancer type ambiguous, and making a prognosis manually based on every single region of it can be challenging. Those new techniques can efficiently highlight regions associated with a certain cancer type. Finally, we have shown for the first time the potential to use deep-learning on histopathology images to predict some gene mutations at an early stage.

## Methods

The overall steps described in this section are summarized in **Figure 1** and described in the following sections.

### Dataset of 1,634 whole-slide images

Our dataset comes from the NCI Genomic Data Commons^31^ which provides the research community with an online platform for uploading, searching, viewing and downloading cancer-related data. All freely available slide images of Lung cancer were uploaded from this source. We studied the automatic classification of “solid tissue normal” and “primary tumor” slides using a set of respectively 459 and 1175 eosin stained histopathology whole-slide images. Then, the “primary tumor” were classified between LUAD and LUSC types using a set of respectively 567 and 608 of those whole-slide images.

### Image pre-processing generates 987,931 tiles

The slides were tiled in non-overlapping 512x512 pixel windows at a magnification of x20 using the openslide library^32^ (533 of the 2167 slides initially uploaded were removed because of compatibility and readability issues at this stage). The slides with a low amount of information were removed, that is all the tiles where more than 50% of the surface is covered by background (where all the values are below 220 in the RGB color space). This process generated nearly 1,000,000 tiles.

**Table 2.**
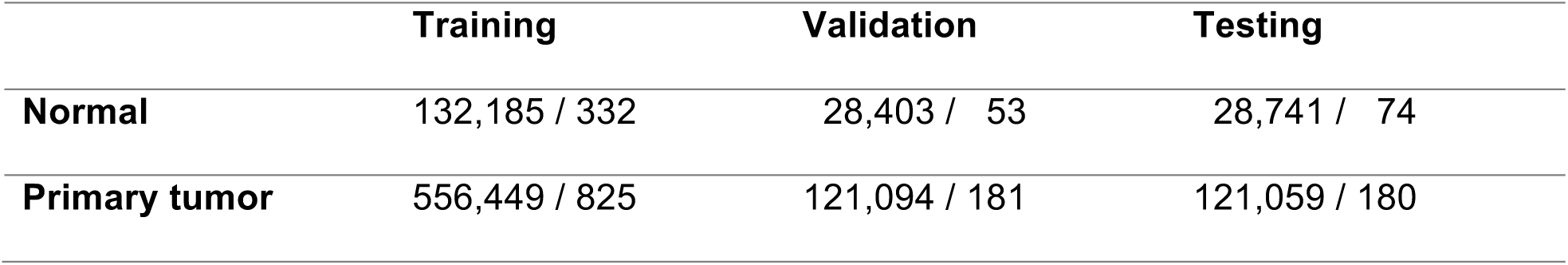
Dataset information for normal vs tumor classification: number of tiles / slides in each category.

**Table 3.** Dataset information for LUAD vs LUSC classification: number of tiles / slides in each category.

|  | Training | Validation | Testing |
| --- | --- | --- | --- |
| **LUAD** | 255,975 / 403 | 55,721 / 85 | 55,210 / 79 |
| **LUSC** | 300,474 / 422 | 65,373 / 96 | 65,849 / 91 |

### Deep learning with Convolutional Neural Network

We used 70% of those tiles for training, 15% for validation, and 15% for final testing (**Table 2** and **Table 3**). The tiles associated with a given slide were not separated but associated as a whole to one of these sets to prevent overlaps between the three sets. Typical CNN consist of several levels of convolution filters, pooling layers and fully connected layers. We based our model on inception v3 architecture^19^. This architecture makes use of inception modules which are made of a variety of convolutions having different kernel sizes and a max pooling layer. The initial 5 convolution nodes are combined with 2 max pooling operations and followed by 11 stacks of inception modules. The architecture ends with a fully connected and then a softmax output layer. For “normal” vs “tumor” tiles classification, we fully trained the entire network. For the classification of type of cancer, we followed and compared different approaches to achieve the classification: transfer learning, which includes training only the last fully-connected layer, and training the whole network. Tests were implemented using the Tensorflow library (tensorflow.org).

### Transfer learning on inception v3

We initialized our network parameters to the best parameter set that was achieved on ImageNet competition. We then fine-tuned the parameters of the last layer of the network on our data via back propagation. The loss function was defined as the cross entropy between predicted probability and the true class labels, and we used RMSProp^33^ optimization, with learning rate of 0.1, weight decay of 0.9, momentum of 0.9, and epsilon of 1.0 method for training the weights. This strategy was tested for the binary classification of LUAD vs LUSC.

### Training the entire inception v3 network

The inception v3 architecture was fully trained using our training datasets and following the procedure described in ^34^. Similar to transfer learning, we used back-propagation, cross entropy loss, and RMSProp optimization method, and we used the same hyperparameters as the transfer learning case, for the training. In this approach, instead of only optimizing the weights of the fully connected layer, we also optimized the parameters of previous layers, including all the convolution filters of all layers. This strategy was tested on three classifications: normal vs tumor, LUAD vs LUSC and Normal vs LUAD vs LUSC. The training jobs were run for 500,000 iterations. We computed the cross-entropy loss function on train and validation dataset, and used the model with best validation score as our final model. We did not tune the number of layers or hyper-parameters of the inception network such as size of filters.

### Identification of gene mutations

To study the prediction of gene mutations from histopathology images, we modified the inception v3 to perform multi-task classification rather than a single task classification. Each mutation classification was treated as a binary classification, and our formulation allowed multiple mutations to be assigned to a single tile. We optimized the average of the cross entropy of each individual classifier. To implement this method, we replaced the final softmax layer of the network with a sigmoid layer, to allow each sample to be associated with several binary labels ^35^. We used RMSProp algorithm for the optimization, and fully trained this network for 500k iterations using only LUAD whole-slide images, each one associated with a 10-cell vector, each cell associated to a mutation and set to 1 or 0 depending on the presence or absence of the mutation. Only the most common mutations were used (**Table 4**), leading to a training set of 223,185 tiles. Training and validation were done over 500,000 iterations (**Figure supp 3**). The test was then achieved on the tiles, and aggregation on the 62 test-slides where at least one of these mutations is present was done only if the tile was previously classified as “LUAD” by the Normal/LUAD/LUSC 3-classes classifier.

**Table 4.** Gene included in the multi-output classification and the percentage of patients with LUAD in the database where the genes are mutated.

| Gene<br>mutated | TP53 | LRP1B | KRAS | KEAP1 | FAT4 | STK11 | EGFR | FAT1 | NF1 | SETBP1 |
| --- | --- | --- | --- | --- | --- | --- | --- | --- | --- | --- |
| %Patients | 50 | 34 | 28 | 18 | 16 | 15 | 12 | 11 | 11 | 11 |

### Results analysis

Once the training phase was finished, the performance was evaluated using the testing dataset which is composed of tiles from slides not used during the training. We then aggregated the probabilities for each slide using two methods: either average of the probabilities of the corresponding tiles, or percentage of tiles positively classified. The ROC (Receiver Operating Characteristic) curves and the corresponding AUC (Area Under the Curve) were computed in each case. Tumor slides could contain a certain amount of “normal” tiles. Therefore, we also checked how the ROC & AUC were affected when tiles classified as “normal” were removed from the aggregation. Heat-maps were also generated for some tested slide to visualize the differences between the two approaches and identify the regions associated with a certain cancer type. To visualize the regions of a given tile which were important for the algorithm to take a decision, a rolling mask was applied to the tile. The masked tile was then fed to the network to analyze how the classification is affected. 128x128 pixel overlapping masks were generated over the whole tile with 87.5% overlapping between adjacent masks.

## Acknowledgments

This work has utilized computing resources at the High-Performance Computing Facility at NYU Langone Medical Center. The slide images and the corresponding cancer information were uploaded from the Genomic Data Commons portal (https://gdc-portal.nci.nih.gov) and are in Whole or part based upon data generated by the TCGA Research Network (http://cancergenome.nih.gov/).The data used were publicly available without restriction, authentication or authorization necessary.

